# Novel hydrogen- and iron-oxidizing sheath-producing Zetaproteobacteria thrive at the Fåvne deep-sea hydrothermal vent field

**DOI:** 10.1101/2023.06.20.545787

**Authors:** Petra Hribovšek, Emily Olesin Denny, Håkon Dahle, Achim Mall, Thomas Øfstegaard Viflot, Chanakan Boonnawa, Eoghan P. Reeves, Ida Helene Steen, Runar Stokke

## Abstract

Iron oxidizing Zetaproteobacteria are well-known to colonize deep-sea hydrothermal vent fields around the world where iron-rich fluids are discharged into oxic seawater. How inter-field and intra-field differences in geochemistry influence the diversity of Zetaproteobacteria, however, remains largely unknown. Here, we characterize Zetaproteobacteria phylogenomic diversity, metabolic potential, and morphologies of the iron oxides they form, with a focus on the recently discovered Fåvne vent field. Located along the Mohns ridge in the Arctic, this vent field is a unique study site with vent fluids containing both iron and hydrogen with thick iron microbial mats (Fe mats) covering porously venting high-temperature (227-267 °C) black smoker chimneys. Through genome-resolved metagenomics and microscopy, we demonstrate that the Fe mats at Fåvne are dominated by tubular iron oxide sheaths, likely produced by Zetaproteobacteria of genus *Ghiorsea*. With these structures, *Ghiorsea* may provide a surface area for members of other abundant taxa such as Campylobacterota, Gammaproteobacteria and Alphaproteobacteria. Furthermore, *Ghiorsea* likely oxidizes both iron and hydrogen present in the fluids, with several *Ghiorsea* populations co-existing in the same niche. Homologues of Zetaproteobacteria Ni,Fe hydrogenases and iron oxidation gene *cyc2* were found in genomes of other community members, suggesting exchange of these genes could have happened in similar environments. Our study provides new insights into Zetaproteobacteria in hydrothermal vents, their diversity, energy metabolism and niche formation.

**Importance:** Knowledge on microbial iron oxidation is important for understanding the cycling of iron, carbon, nitrogen, nutrients, and metals. The current study yields important insights into the niche sharing, diversification, and Fe(III) oxyhydroxide morphology of *Ghiorsea*, an iron- and hydrogen oxidizing Zetaproteobacteria representative belonging to ZetaOTU9. The study proposes that *Ghiorsea* exhibits a more extensive morphology of Fe(III) oxyhydroxide than previously observed. Overall, the results increase our knowledge on potential drivers of Zetaproteobacteria diversity in iron microbial mats and can eventually be used to develop strategies for the cultivation of sheath-forming Zetaproteobacteria.

## 1 Introduction

Chemolithoautotrophic iron-oxidizing (Fe-oxidizing) bacteria are frequently observed at deep-sea hydrothermal vents associated with Fe(II)-rich fluids. Fe-oxidizing bacteria can obtain energy by oxidation of Fe(II) under microaerobic conditions, thereby competing with abiotic Fe oxidation, or some can grow anaerobically using nitrate as terminal electron acceptor (1–4). Fe-oxidizers influence biogeochemical cycling of iron and other elements through transforming the soluble Fe(II) to insoluble Fe(III), which often takes the form of Fe(III) oxyhydroxides. These Fe(III) oxyhydroxides have the ability to co-precipitate and adsorb carbon, nutrients, and heavy metals (5–10). Fe-oxidizing bacteria can also play a role in corrosion (11, 12) and bioremediation of metal pollution and recovery of resources (13, 14). Seafloor hydrothermal fluids are typically rich in diverse electron donors such as hydrogen sulfide and methane, but with variable hydrogen and Fe contents (15–18). The exact chemical composition of these fluids varies widely between and often within vent fields, depending on the hydrothermal system’s geological setting, and exerts a strong influence on the microbial communities present (19–21).

Fe-oxidizing bacteria are found in iron microbial mats (Fe mats) around the globe, such as at the Kama‘ehuakanaloa (Lō‘ihi) seamount (4, 22–25), Vailuluʻu seamount (26), Mid-Atlantic Ridge (27, 28), the Mariana region (29–34), Kermadec Arc (35), South Tonga Arc (36), at Mid-Cayman Ridge (37), and at Arctic Mid-Ocean Ridges (38–40). The dominant Fe-oxidizing bacteria in these Fe mats are Zetaproteobacteria, first proposed as a class in 2007 (41) and collectively divided into operational taxonomic units specific to subgroups of Zetaproteobacteria (ZetaOTUs) (42). To obtain energy for CO_2_ fixation, Zetaproteobacteria oxidize soluble Fe(II) under low oxygen conditions (1). Micrometer-scale structures composed largely of extracellular polymeric substances and precipitated Fe(III) oxyhydroxides are often formed as a result of their Fe metabolism. The morphology of Fe(III) hydroxides vary between types of Zetaproteobacteria. Some produce twisted stalks, others produce hollow tubular sheaths, bifurcating tubular structures, or dreads (22, 24, 43–45). It has been hypothesized that these structures prevent the cells from becoming encrusted in Fe and that they keep the cells within the gradient of oxygen and Fe required for growth (24, 41). While stalk formation genes have recently been discovered (46, 47), the molecular mechanisms for formation of other structures are not well studied and not all morphologies have a known isolated representative. To date, no sheath-forming Zetaproteobacteria have been cultured, nor have the Zetaproteobacteria responsible for sheath formation been identified. Since sheaths and stalks make up the majority of Fe mats, stalk- and sheath-forming Zetaproteobacteria are recognized as keystone species that engineer the structure of these mats, providing a suitable environment for other species (24).

Only a few Fe oxidizers belonging to the Zetaproteobacteria have been cultured, most of which are members of the genus *Mariprofundus* (41, 43, 48, 49). The *cyc2* gene has been validated as the main gene involved in the Fe oxidation pathway of Zetaproteobacteria and other bacteria in near-neutral pH environments (34, 50–52). While most of Zetaproteobacteria are strict Fe oxidizers, it has been shown that *Ghiorsea bivora*, a ZetaOTU9 representative, can obtain energy from hydrogen oxidation by using hydrogen as either the sole electron donor or in combination with Fe(II) (53). The co-occurrence of Fe(II) and H_2_ may play an important role in defining the niche of ZetaOTU9 (1). However, we have a limited understanding of the functioning of Fe oxidizers that also use H_2_ and how this affects their diversity and ecology. In this paper, we contribute towards narrowing this knowledge gap.

At the recently discovered Fåvne deep-sea hydrothermal vent field located on the Mohn’s Ridge (54–56), dense Fe mats cover porous black smoker chimney surfaces at *in situ* temperatures of ∼50 °C. The venting fluids at Fåvne contain both dissolved H_2_ and Fe(II) as abundant energy sources (55) and thus, Fåvne is a valuable study site to investigate Fe oxidizers that can use H_2_ as an alternate electron donor. Our genome-resolved metagenomics and microscopy study characterizes Fåvne Fe mats as a deep-sea hydrothermal habitat formed using abundant byproducts of novel sheath-forming Fe-oxidizing Zetaproteobacteria that potentially also utilize H_2_. The identification of sheath-producing *Ghiorsea* belonging to ZetaOTU9 extends previous knowledge on Fe(III) oxyhydroxide morphologies. In addition, our results suggest that hydrogen could be the main driver of diversity of Zetaproteobacteria interacting with vent fluids containing both Fe(II) and H_2_, where flexible lithotrophic energy metabolism of *Ghiorsea* provides an advantage.

## 2 Results

### Zetaproteobacteria produce Fe(III) oxyhydroxide tubular sheaths in Fe mats at Fåvne

The porous black smoker chimneys at Fåvne show focused-flow venting at 227 °C of fluids containing abundant Fe(II) and H_2_ (55), and support growth of extensive Fe mats covering tall black smoker chimney spires (Figure 1a, 1b). The temperature within the Fe mats close to the venting orifice on chimney exteriors was measured at ∼50 °C (Figure 1b). The chimney structures appear to lack defined central conduits (54), leading to copious venting of hydrothermal fluids (55) through permeable and porous chimney walls. Analysis of the microbial community composition based on metagenome-assembled genome (MAG) coverage (and supported by 16S rRNA gene read abundance), revealed that Zetaproteobacteria comprise 7% of the community (FeMat, Supplementary Material 4 Table S1). Other lineages frequently observed at vents (57–61) were more abundant in the Fe mat sample than Zetaproteobacteria, including members of Gammaproteobacteria and Campylobacterota (mainly *Sulfurovum*), which comprise 31% and 30% of the community, respectively (Figure S12). A single Alphaproteobacteria *Robiginitomaculum* MAG comprised ∼2% of total MAG coverage. Interestingly, three high-quality (CheckV) viral genomes (vMAGs) identified in the Fe mat are predicted to have abundant Fe mat bacteria *Sulfurimonas* (Campylobacterota) and Gammaproteobacteria as potential hosts (Table S9).

**Figure 1.**
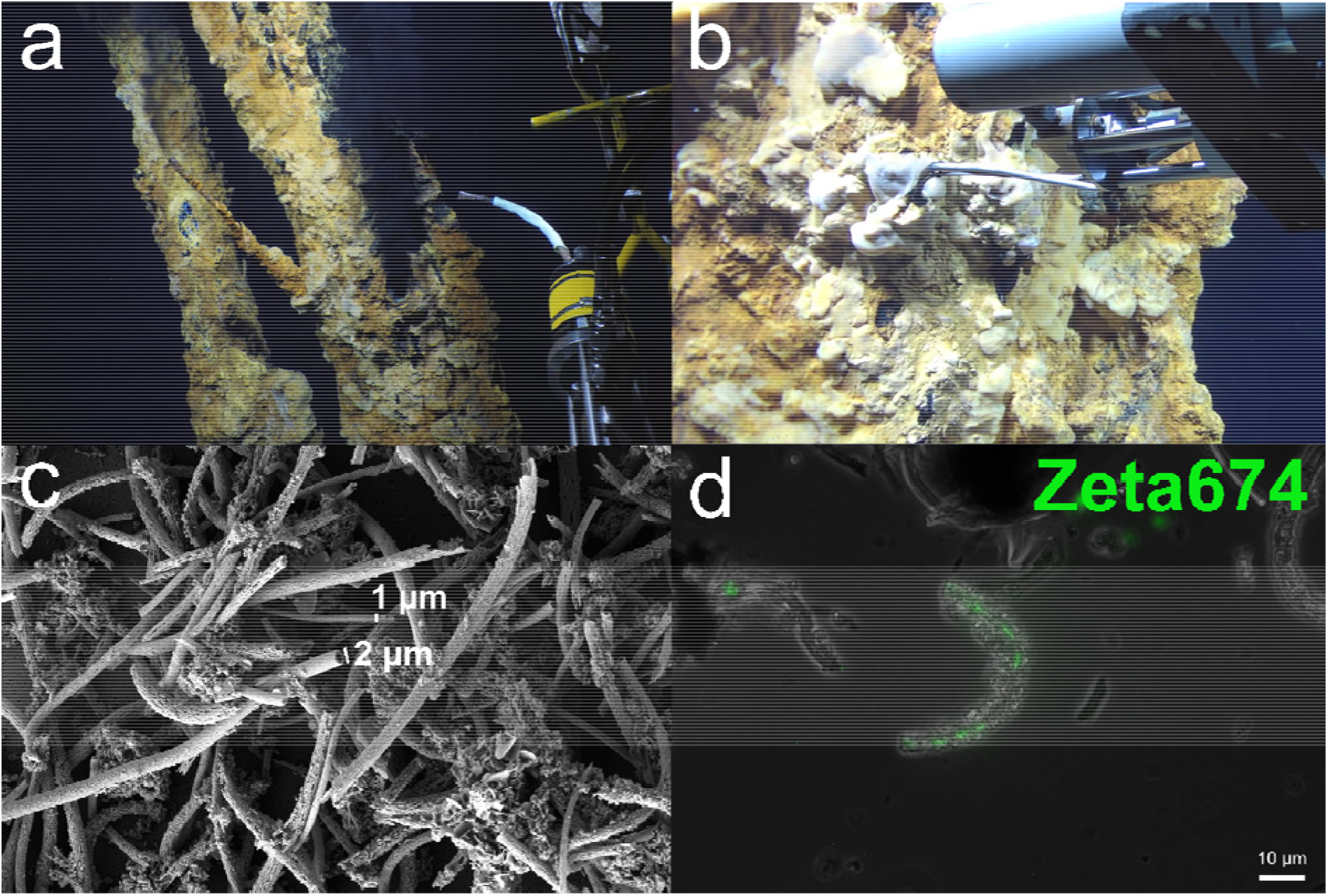
Fe mats at Fåvne are dominated by Fe(III) oxyhydroxide sheaths produced by Zetaproteobacteria. a) Fe mats on black smoker chimneys at Fåvne vent field with hydraulic suction device (biosyringe) used for sampling on the right. b) Measuring temperatures within Fe mats using an Isobaric Gas-Tight sampler (55). c) Fe mats dominated by Fe(III) oxyhydroxide sheaths of two different widths produced by Zetaproteobacteria, scanning electron microscopy. Two Fe(III) oxyhydroxide sheath morphotypes, either 2 μm or 1 μm wide. d) Zetaproteobacteria cells inside Fe(III) oxyhydroxide sheaths (stained with Zeta674 FISH probe). Overlay of phase-contrast and florescence images.

Scanning electron microscopy (SEM) revealed that tubular Fe(III) oxyhydroxide sheaths dominate the black smoker Fe mats (Figure 1c, Figure S2, Figure S4). Fe(III) oxyhydroxide sheaths were the only morphotype observed, the majority of which are about 2 μm wide, with thinner 1 μm sheaths also present, albeit less frequently. The Zeta674 FISH probe detected Zetaproteobacteria cells inside the sheaths (Figure 1d, Figure S3), identifying tubular sheath-forming Zetaproteobacteria as major Fe-oxidizers in the Fe mat.

### Phylogeny of Fe-oxidizing Zetaproteobacteria at Fåvne

Phylogenomics analysis was performed to elucidate the taxonomy of the sheath-forming Zetaproteobacteria community in the Fe mat. Five of the 127 high- and medium-quality MAGs (62) reconstructed from the Fe mat were classified as the Zetaproteobacteria genus *Ghiorsea* (GTDB and 16S rRNA genes). All but one of the *Ghiorsea* MAGs are new species-representative genomes based on publicly available genomes of Zetaproteobacteria and a 95% average nucleotide identity (ANI) cut-off (63, 64). Phylogenomic and average amino acid (AAI) identity analyses among *Ghiorsea* identified two distinct clusters previously not described for the genus (designated Cluster A and B, Figure 2, Figure S11). The two Fåvne MAGs in Cluster A were affiliated with the symbiont *Ghiorsea* from the vent shrimp *Rimicaris* (65), *Ghiorsea* from Mid-Cayman Rise and North Pond (66), whereas the three other Fåvne MAGs in Cluster B were affiliated with the cultivated *G. bivora* (53) and *Ghiorsea* from Urashima (34). The two dominating *Ghiorsea* Fåvne MAGs, Faavne_M6_B18 and AMOR20_M1306, are members of each of these clusters, and were present at 5 and 2%, respectively. Based on ANI estimates, the closest publicly available genome to the highest quality *Ghiorsea* MAG from Fåvne is a *Ghiorsea* MAG (64.8% complete) from a cold oxic subseafloor aquifer (66) with an ANI value of 81.3% (Figure 10, Figure S11). The same Fåvne MAG has an ANI value of 77.7% to *Ghiorsea bivora* (53), an isolated Zetaproteobacteria ZetaOTU9 representative, while the 16S rRNA gene has 96.8% identity to the isolate sequence.

**Figure 2.**
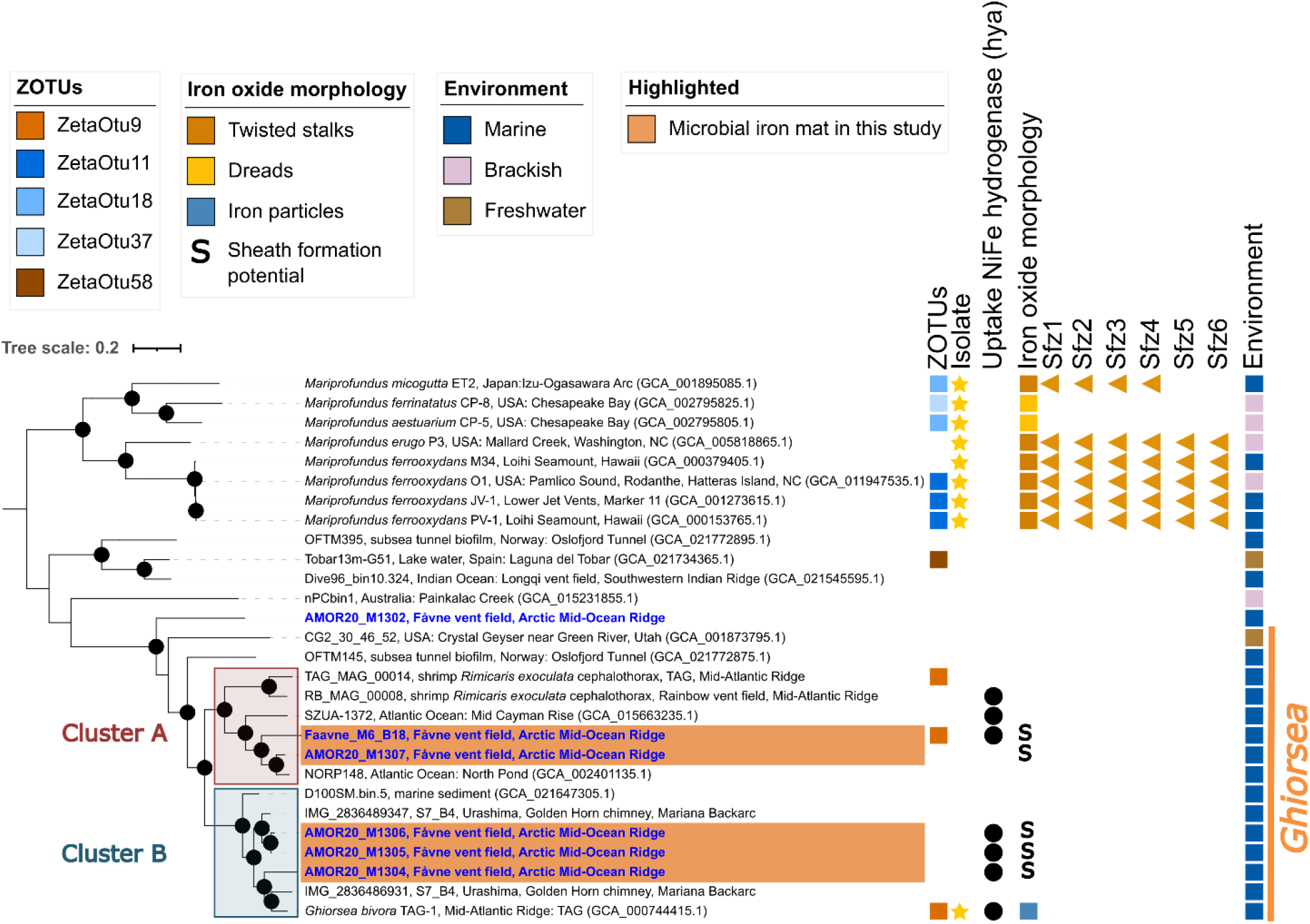
Phylogeny of Zetaproteobacteria in black smoker Fe mats at Fåvne. The tree is based on a concatenated alignment of a manually curated set of 12 single copy gene markers (Table S5) using MAGs from this study and references. Blue genomes have been reconstructed from the Fåvne vent field. Genomes highlighted in orange are present in Fe mat associated to a focused-flow black smoker (Ghiorsea, ZetaOTU9). MAGs above 0.5 coverage were selected. Sfz1-6: potential stalk formation genes in Zetaproteobacteria. Black node circles mark branches with support values higher than 75% with standard bootstrapping and 1000 iterations. The maximum likelihood tree with substitution model Qpfam+F+I+I+R7. The Ghiorsea genus is based on GTDB taxonomy r202 and AAI values within the proposed 65% AAI cut-off for genus (133).

To compare the taxonomy of Zetaproteobacteria in Fe mats with those present at other locations at Fåvne, we recovered MAGs from other Fåvne sampling sites (Supplementary Material 4 Table S1). While black smoker Fe mats Zetaproteobacteria were all assigned to the genus *Ghiorsea*, low-temperature diffuse venting at Fåvne supported a higher number of other Zetaproteobacteria taxa (Table S11). 28 unique species-representative genomes of Zetaproteobacteria were recovered at Fåvne (based on 95% ANI cutoff and publicly available MAGs) consisting of high- and medium-quality MAGs (average completeness 81.7%, contamination 2.1% based on CheckM2; Table S4). These Fåvne MAGs were associated with 2 families defined by GTDB and 7 defined genera, with 3 MAGs remaining unclassified to genus level and most of taxa lacking cultured representatives (Figure S9).

### *Ghiorsea* in Fåvne Fe mats can oxidize H_2_ in addition to Fe(II)

To assess the physiology of *Ghiorsea* species in the Fe mat, annotated genomes were analyzed for potential electron donors and acceptors. In alignment with the presence of H_2_ and Fe(II) in endmember fluids at Fåvne (55), genes encoding all subunits of a transmembrane H_2_-uptake Ni,Fe hydrogenase (Group 1d) and Cyc2 for Fe(II) oxidation were identified in Fåvne *Ghiorsea* genomes belonging to both cluster A and B (Figure 2, Figure 4). In addition, codon usage bias analysis predicts high expression of Fe(II) oxidation and H_2_ oxidation genes of the *Ghiorsea* MAGs from Fåvne (Table S12). A broader functional screening revealed that H_2_-based metabolism with a Group 1d hydrogenase is common to other dominant MAGs within the Fe mat belonging to the Gammaproteobacteria, Ignavibacteria, Calditrichia, KSB1 and Aquificae (Figure S18, Table S8A). *Ghiorsea* and some Gammaproteobacteria in the Fe mat also encode genes of a Ni,Fe H_2_-sensing hydrogenase histidine kinase-linked Group 2b (*hup*) located in the cytosol responsible for activating hydrogenase expression (Figure S15) (67). In contrast, hydrogenases were not detected in Zetaproteobacteria MAGs not belonging to genus *Ghiorsea* from other locations at Fåvne.

**Figure 3.**
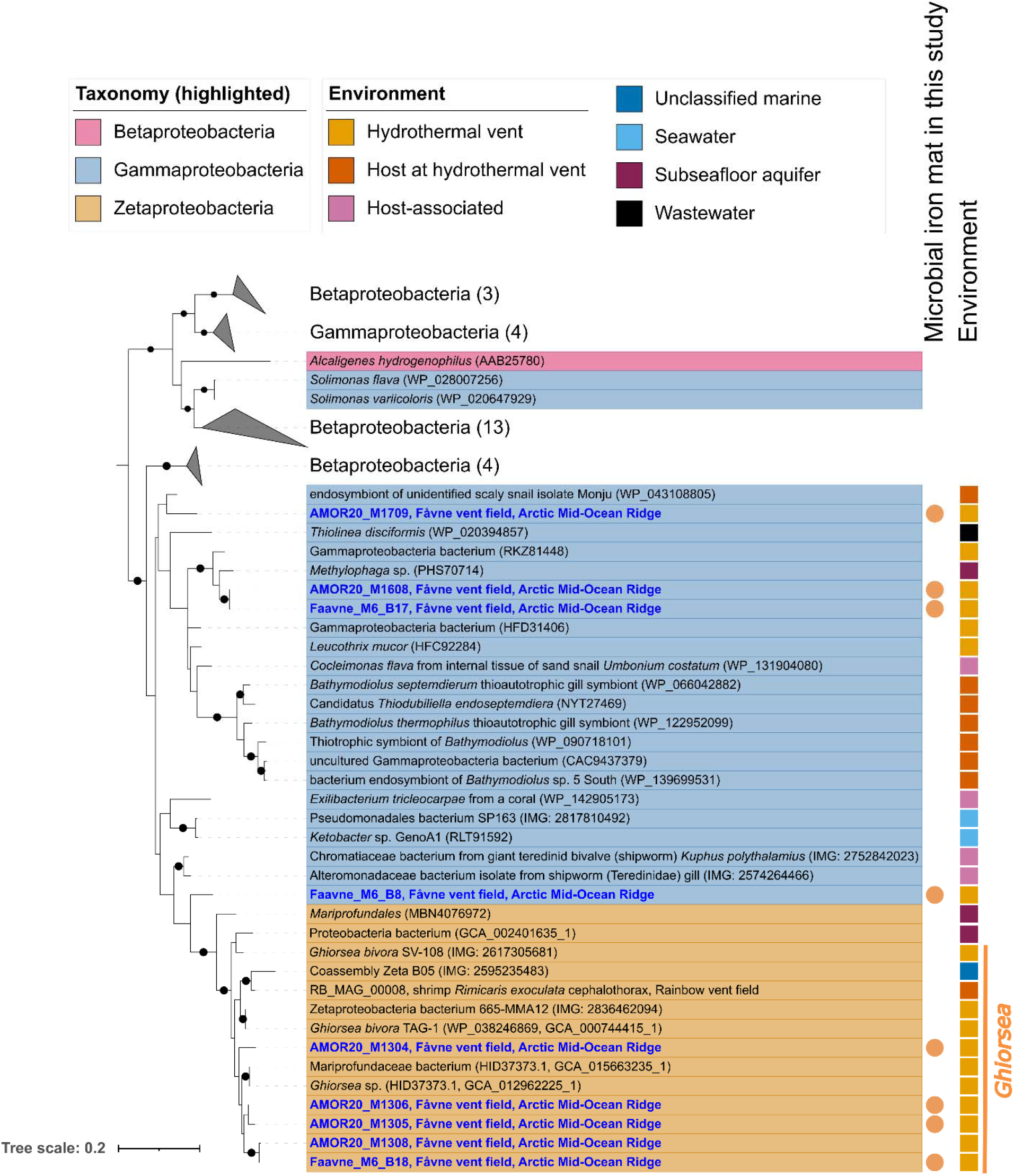
Phylogeny of the large subunit of uptake NiFe hydrogenase (hya; 1d). Phylogenetic tree of the large subunit of uptake NiFe hydrogenase (hya; 1d) present in MAGs in the black smoker Fe mat and in all publicly available Zetaproteobacteria genomes, with closest relative reference using BLAST. Blue MAGs have been reconstructed from the Fåvne vent field. Black node circles mark branches with support values higher than 75% with standard bootstrapping and 1000 iterations. Maximum likelihood tree with substitution model LG+I+I+R7.

**Figure 4.**
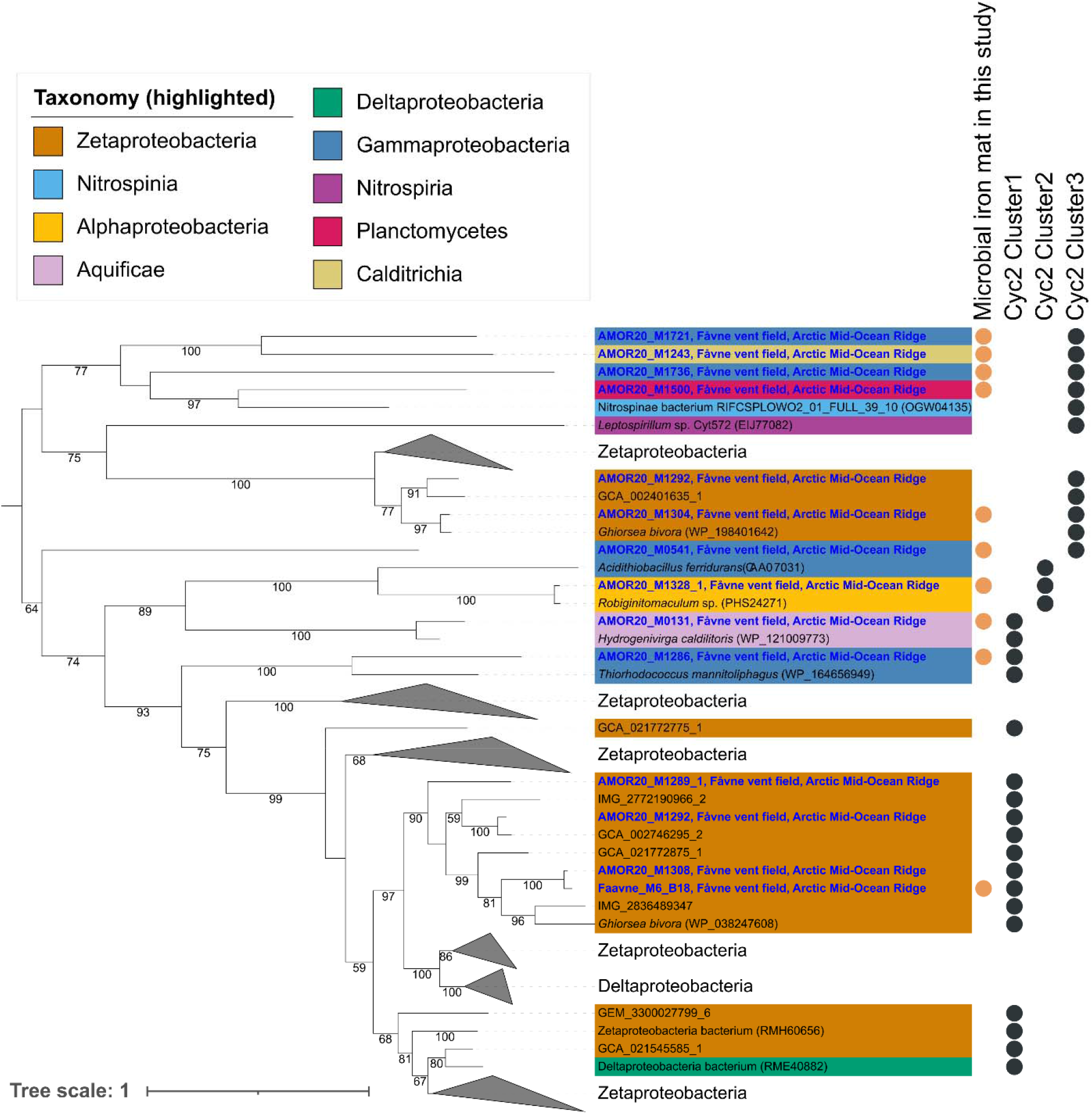
Phylogeny of outer membrane cytochrome Cyc2. Phylogenetic tree of Fe oxidation cytochrome Cyc2 present in MAGs in the black smoker Fe mat including amino acid sequences from all publicly available Zetaproteobacteria genomes with closest relative references using BLAST. Blue labels are sequences from MAGs reconstructed from the Fåvne vent field in the current study. Support values for branches are calculated with standard bootstrapping and 1000 iterations. The maximum likelihood tree was constructed using the substitution model Qpfam+F+I+I+R5.

A phylogenetic tree constructed using the large subunit of the transmembrane Ni,Fe hydrogenase (Figure 3, Figure S16) to assess the evolutionary relationships of encoded hydrogenases, reveals the close relationship of Fåvne *Ghiorsea* hydrogenases to hydrogenases of other *Ghiorsea* (53, 65) and Gammaproteobacteria. Interestingly, the closest non-Zetaproteobacteria homologue was identified as a hydrogenase from a Gammaproteobacteria MAG (o. Granulosicoccales, encoding genes for sulfur oxidation) from the same Fe mat.

The *cyc2* gene has previously been identified as one of the key genes in Fe(II) oxidation, with three distinct phylogenetic clusters of functionally verified and biochemically characterized representative Fe oxidases (51, 68, 69). Identified Cluster 3 and Cluster 1 *cyc2* genes of *Ghiorsea* in the Fe mat at Fåvne showed highest similarity to *cyc2* in other *Ghiorsea* MAGs from hydrothermal vents. Cyc2 genes were also identified in MAGs belonging to Gammaproteobacteria, Alphaproteobacteria, Aquificae, Planctomycetes and Calditrichia (Figure 4, Table S8B).

Manganese and iron are well known to co-vary in high-temperature black smoker fluids (70). Gene annotations revealed the presence of several genes in Gammaproteobacteria putatively involved in manganese oxidation, such as *mcoA, mopA* and *moxA* (71–75). Furthermore, preliminary proteomics analysis of abundant proteins in black smoker Fe mats show expressed McoA (Table S10, Supplementary Material 1 Text2).

### Metabolism of the microbial community in Fe mats

Analysis of the genomic content of the 25 most abundant MAGs (Table S3), contributing to 87% of the binned coverages, identified genes for the oxidation of sulfur compounds, H_2_, CH_4_, and NH_4_^+^ (Figure 5, Figure S14). Terminal oxidases found in *Ghiorsea* MAGs were cbb3-type cytochrome c oxidases, indicating an adaptation to low oxygen concentrations (76). Other MAGs contained both cbb3-, aa3-type cytochrome c oxidases, and cytochrome bd-I ubiquinol oxidases. Nitrate and nitrite reductase genes were identified in *Ghiorsea* MAGs, indicating a possibility of an auxiliary anaerobic metabolism (23, 77). Arsenate reductase was also detected.

**Figure 5.**
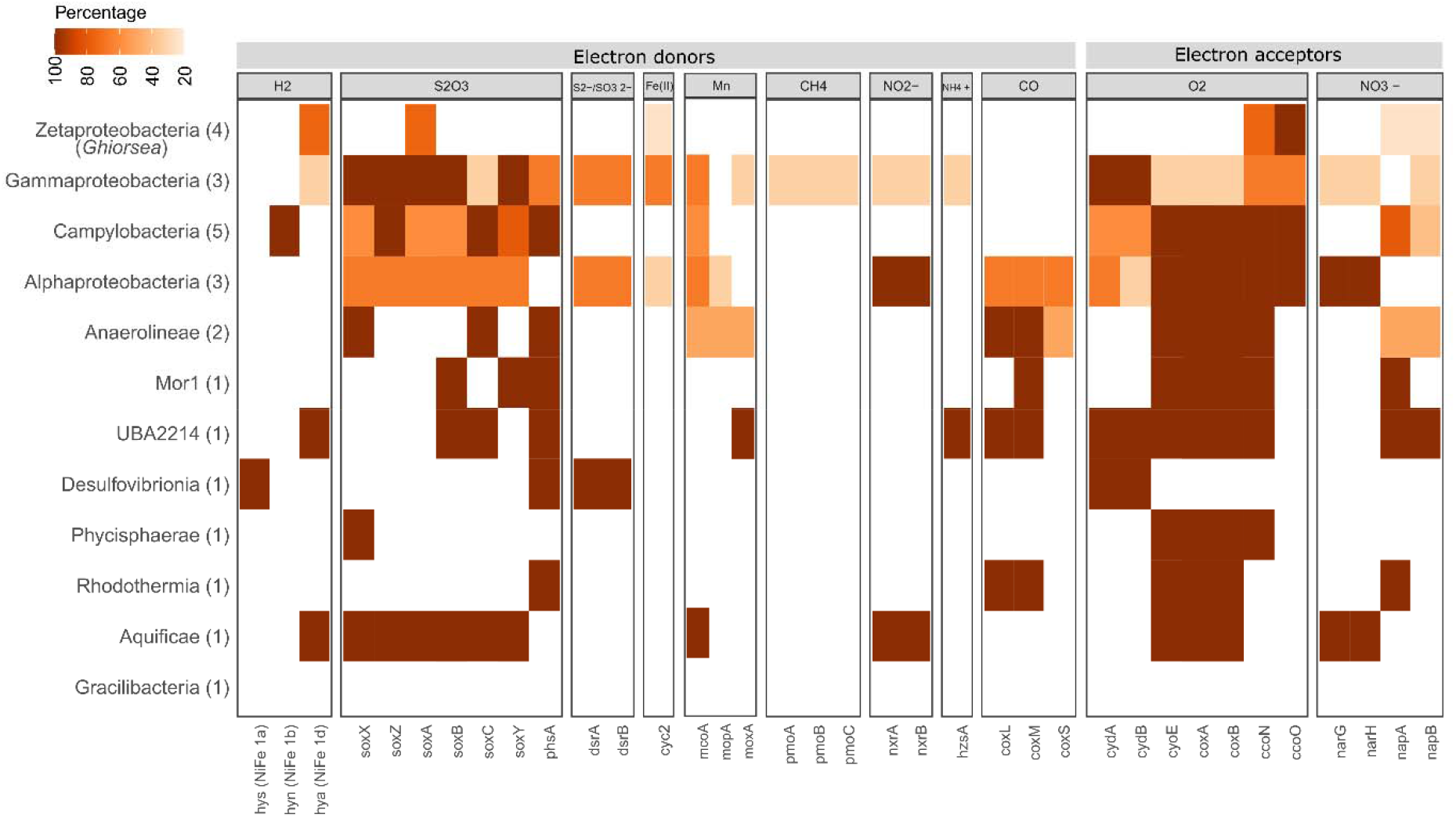
Functional characterization of the top 25 abundant MAGs in the black smoker Fe mat. Distribution of genes involved in the utilization of a range of electron donors and electron acceptors. The number of genomes in each taxonomic class cluster is indicated in parenthesis and the color gradient refers to the percentage of genomes per class that encode the genes. Top 25 most abundant MAGs account for 87% of MAG coverages.

Enzymes involved in carbon fixation were identified (Figure S13) for expected CO_2_ fixation pathways in representative lineages (1, 78–80), however key genes for the serine variant of the reductive glycine pathway were observed in a Campylobacterota MAG (81).

Given that Fe(III) oxyhydroxides adsorb heavy metals (8), an analysis of heavy metal resistance genes within the full metagenomic assembly of the Fe mat was performed. Heavy-metal resistance genes were identified for copper, cobalt, sodium acetate, chromium, tellurium, selenium and silver (Table S7).

## 3 Discussion

Fe-oxidizing Zetaproteobacteria are globally distributed, yet our knowledge on the importance of hydrogen for their distribution is still limited. Here, we phylogenetically and functionally characterized Fe(II)- and H_2_-oxidizing Zetaproteobacteria from the Fåvne vent field belonging to *Ghiorsea* genus, adding four novel species-representative genomes predicted to use H_2_. We reconstructed 28 novel species-representative genomes of diverse Zetaproteobacteria taxa, extending the known Zetaproteobacteria diversity. Based on Zetaproteobacteria distribution at Fåvne and encoded uptake hydrogenases, we demonstrate that H_2_ availability indeed plays a role in the niche diversity of Zetaproteobacteria. Multiple species of *Ghiorsea* share the H_2_ oxidation capacity in Fe mats at Fåvne, possibly sharing one niche.

Until recently, the identity of sheath-forming Zetaproteobacteria has remained elusive. We show that at least two populations of *Ghiorsea* (ZetaOTU9) produce Fe(III) oxyhydroxide sheaths and form dense Fe mats.

### Hydrogen as a driver of Zetaproteobacteria diversification

Most Zetaproteobacteria genera are metabolic specialists only able to obtain energy from the oxidation of Fe(II). The only known cultivated exception is *G. bivora,* capable of using H_2_ simultaneously with Fe(II), or as sole electron donor (53). It has been suggested that members of *Ghiorsea* (ZetaOTU9) only occupy environments rich in Fe(II), but also combined with predicted presence of H_2_, such as at hydrothermal vents (53), in corrosion of steel (12, 82) and mineral weathering (1, 82, 83). The presence of hydrogen in these *Ghiorsea* environments has mainly been based on hypotheses until the current study. Remarkably, in the Fe mat close to the venting orifice at Fåvne and in contact with fluids containing abundant H_2_, the reconstructed Zetaproteobacteria MAGs are represented by only *Ghiorsea* within ZetaOTU9.

In contrast, a higher diversity of Zetaproteobacteria is present in low-temperature diffuse-venting areas at around ∼10 °C (Table S11, Figure S7, Figure S8). All these genomes, except for *Ghiorsea*, lack uptake hydrogenases (Figure S9). Low-temperature diffuse-venting areas may reflect a low availability of H_2_ relative to Fe(II), lost by abiotic or other subsurface mixing processes and low temperature fluid formation (84). *Ghiorsea*, with its hydrogen uptake capability, emerges as the sole specialist in the presence of H_2_. While *Ghiorsea* is also observed in likely H_2_-poor diffuse-flow environments, the higher diversity of Zetaproteobacteria, reflected by a diversity of Fe(III) oxyhydroxide structures (Figure S5, Figure S6), indicates an absence of a monopolizing niche player. This pattern of distribution supports the hypothesis that H_2_ acts as a niche-determining factor for *Ghiorsea* at Fe(II)-rich hydrothermal vents (1). The ability of *Ghiorsea* to utilize H_2_ affords it a competitive advantage as H_2_ is a thermodynamically more favorable energy source than Fe(II), supporting faster cell growth (53). The competitive advantage of growing on H_2_ is likely linked to evading the need for reverse electron flow to replenish the reducing agent NADH needed for CO_2_ fixation (Figure 6).

**Figure 6.**
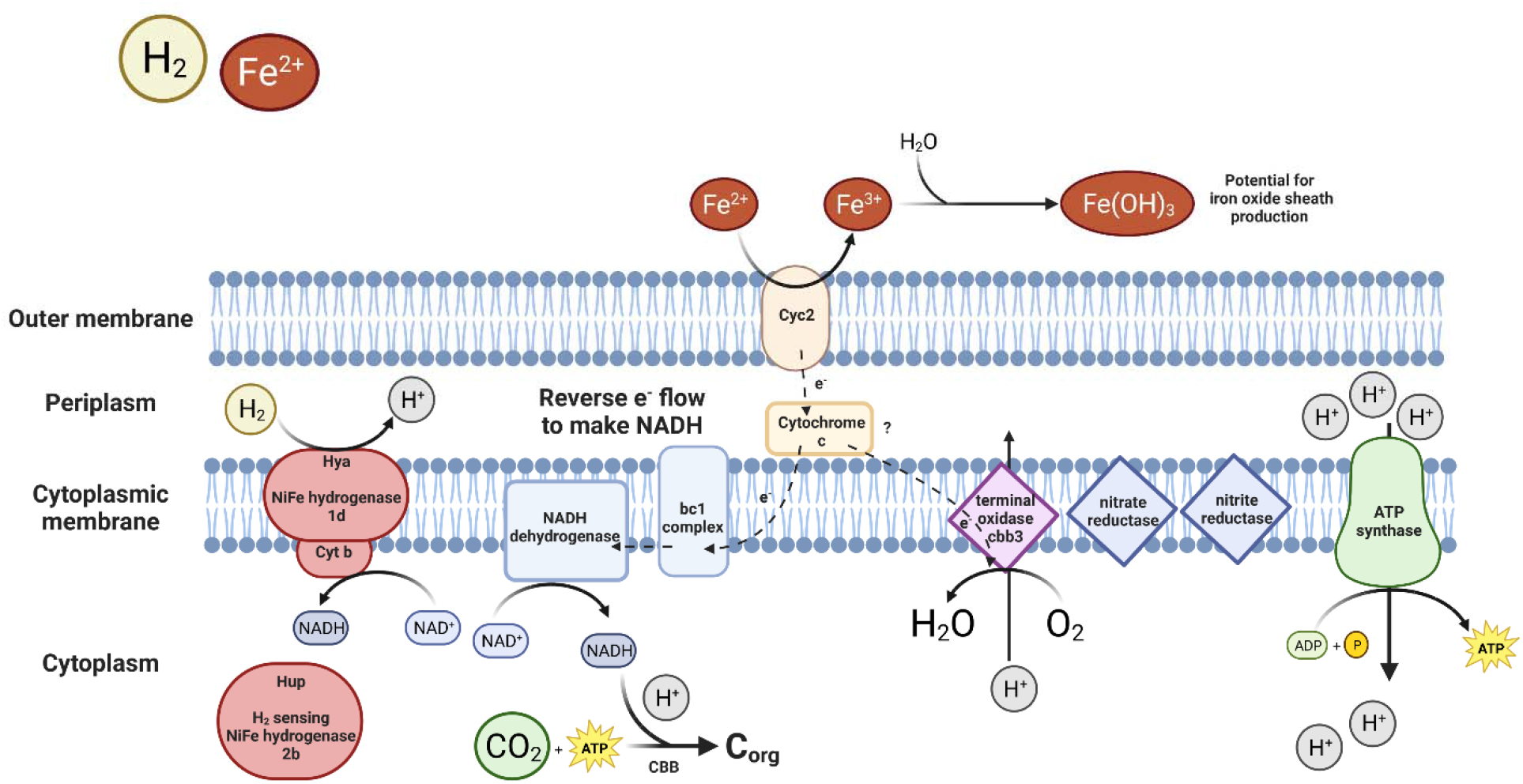
Membrane complexes in Ghiorsea. Electrons coming from the oxidation of Fe^2+^ is passed all the way to the high oxygen affinity terminal oxidase, leading to the generation of a proton motive force. Reverse electron transport is necessary to regenerate NADH needed for CO_2_ fixation. NADH could also get replenished with the help of NiFe uptake hydrogenase instead of the energy-intensive reverse electron transport. Hydrogenase can also donate electrons to the electron transport chain. ATP is generated by ATP synthase. A schematic representation of the metabolic potential of Ghiorsea, based on Ghiorsea MAGs from Fåvne and previous studies (1, 53). Created with BioRender.com.

Hydrogenases restricted to *Ghiorsea* ZetaOTU9 at Fåvne show that potential for growth on H_2_ is a trait limited to ZetaOTU9. Through the analysis of publicly available genomes of Zetaproteobacteria, however, transmembrane uptake hydrogenases were detected in two other species-representative genomes of Zetaproteobacteria outside of *Ghiorsea,* sourced from freshwater and a subseafloor aquifer (Figure S9, Figure S16). Even so, all *Ghiorsea* do not necessarily share the ability to oxidize H_2_. Outside of *Ghiorsea* Clusters A and B, two species-representatives of *Ghiorsea* from freshwater and a subsea tunnel do not appear to possess hydrogenases (Figure 2). Thus far evidence suggests that presence of hydrogenases within *Ghiorsea* may be unique to hydrothermal vents.

Other H_2_ oxidizers besides *Ghiorsea* are also present in the black smoker Fe mat which possess different hydrogenases (Figure 5). Ni,Fe hydrogenases found in *Ghiorsea* MAGs were most closely related to hydrogenase subunits from other *Ghiorsea* and a Gammaproteobacteria MAG within the same Fe mat (Figure 3) and symbiont chemolithotrophic sulfur-oxidizing microorganisms in hydrothermal vent fauna. These observations further strengthen the possibility of horizontal gene transfer of H_2_ oxidation genes between Zetaproteobacteria and lithotrophic sulfur-oxidizing Gammaproteobacteria (53). We hypothesize this may have happened at hydrothermal vents.

Fe mats at Fåvne cover black smokers at temperatures of up to 50 °C (the maximum measured inside of a single Fe mat), which is at the high end of the temperature spectrum where Fe mats and ZetaOTU9 have been observed (1, 53). The role of temperature in distribution patterns of Zetaproteobacteria cannot be ignored, however, minor differences in predicted optimum growth temperatures (Figure S9) indicate fluid composition plays a larger role on the *Ghiorsea* (ZetaOTU9) niche differentiation than temperature. It is worth noting that the Fe mat studied is a bulk sample, where different microenvironments likely exist with varying degrees of exposure to the high-temperature reducing venting fluids. Obtaining small-scale samples and corresponding *in situ* measurements to capture this variability in such environments remains a challenge.

### Several *Ghiorsea* populations at Fåvne share the same metabolic niche

Within the Fe mats at Fåvne, four novel species-representative genomes of *Ghiorsea* (ZetaOTU9) co-exist, belonging to two distinct phylogenetic clusters (Cluster A and Cluster B; Figure 2). Co-existence of multiple *Ghiorsea* populations has also been observed in *Rimicaris* vent shrimp, where the difference in the presence of hydrogenase in *Ghiorsea* MAGs has been proposed to contribute to niche partitioning, avoiding potential competition (65). In contrast, at Fåvne all species-representative *Ghiorsea* genomes seem to possess common metabolic functions including presence of uptake hydrogenases, suggesting that multiple *Ghiorsea* species occupy the same ecological niche at Fåvne. It remains unknown what kind of interactions arise from co-occupying the niche in Fe mats. Possible competitive relationships among closely related populations of Zetaproteobacteria could be responsible for differential distribution across physical space ultimately leading to divergence within the ZetaOTU, as hypothesized for cosmopolitan ZetaOTU2 (85). In this context, the use of genome-resolved metagenomics offers valuable information about distinct subpopulations belonging to the same ZetaOTU and their genomic makeup.

### Production of Fe(III) oxyhydroxide sheaths by members of the Fe- and H_2_-oxidizing genus *Ghiorsea* (Zetaproteobacteria)

In contrast to stalk-forming Zetaproteobacteria (46), the identity of sheath-forming Zetaproteobacteria has not been established through either cultivation or identification of the environmental and genetic drivers for sheath formation. Previous studies have suggested ZetaOTU6, ZetaOTU9 and ZetaOTU15 as candidates for sheath-forming Zetaproteobacteria (25, 27, 82). *Ghiorsea* within ZetaOTU9 accounting for 100% of all Zetaproteobacteria present in the Fe mat and the abundant homogenous Fe(III) oxyhydroxide tubular sheaths containing Zetaproteobacteria cells (Figure 1d, Figure S6, Figure S7) strongly suggest that at Fåvne *Ghiorsea* ZetaOTU9 is uniquely forming these structures. The presence of two size morphotypes of Fe(III) oxyhydroxide sheaths suggests that more than one *Ghiorsea* population is producing Fe(III) oxyhydroxides sheaths. The comparison of MAG relative abundances with abundance of the two sheath variants (Figure 1c) indicates the large 2 μm sheaths are produced by the Faavne_M6_B18 (Cluster A) *Ghiorsea* population while the 1 μm wide sheaths are produced by the AMOR20_M1306 (Cluster B) *Ghiorsea* population. Variable width Fe(III) oxyhydroxide sheaths hypothesized to be created by two different unidentified Zetaproteobacteria has previously been observed in Fe mats at Beebe’s vents (37). Despite these concurring observations, the possibility remains that variation in sheath width is instead a consequence of a later, secondary colonization of the same species under different conditions, resulting in variations in cell size.

Similar to stalk formation, the genetic features for dread-forming Zetaproteobacteria are associated with the presence of distant homologs of stalk-forming Zetaproteobacteria *sfz* genes (46, 47). However, no homologs of the *sfz* genes were identified in *Ghiorsea* MAGs from Fåvne (Sfz1-6 genes; Figure 2) or the full metagenome assembly even at low sequence identity, suggesting a different genetic mechanism for sheath formation.

Whether sheaths are uniquely formed by *Ghiorsea* ZetaOTU9 globally remains an open question. Previous microscopy studies of *Ghiorsea* did not reveal any sheath formation (53, 65), with the cultured representative strains of *Ghiorsea* instead producing amorphous Fe(III) oxyhydroxides particulates during growth on FeCl_2_ (53). This aligns with the observation that closely related species vary in their capacity to produce distinct Fe(III) oxyhydroxide structures (24). Sheath-dominated Fe mat communities have been observed at several locations (22, 24, 26, 27, 35, 37). Given that ZetaOTU9 has been described as having a cosmopolitan distribution (25, 38, 86), member species could be producing Fe(III) oxyhydroxide sheaths in numerous environments worldwide. This emphasizes the importance of microscopy in microbial ecology as not everything can be easily observed through genetic analyses. As preserved biogenic Fe(III) oxyhydroxide structures can help us understand the environmental conditions of early Earth through studying ancient iron oxide deposits (87–90) and can also be potentially used as biosignatures (24, 91), knowledge of a sheath-forming Zetaproteobacteria capable of oxidizing both Fe(II) and H_2_ might prove valuable for the interpretation.

### *Ghiorsea* (ZetaOTU9) is the architect of Fe mats with abundant H_2_

Although sheath-forming *Ghiorsea* is not the most abundant community member (7% relative abundance), it can be considered the main community engineer with respect to the amount of produced material (Figure 7), in agreement with Zetaproteobacteria previously characterized as keystone species and primary colonizers in Fe mats (24, 92). The generation of the architectural character of the Fe mat is subsequently followed by recruitment of other community members (44). At Fåvne, the dense Fe mats close to the venting orifice show a high abundance of Campylobacterota (formerly Epsilonproteobacteria), Gammaproteobacteria, and Alphaproteobacteria; taxa commonly seen in Fe mats (30). Diversity of primary producers in Fe mats at Fåvne appears high in comparison with other hydrothermal vent mats dominated by sulfur oxidizers (58–60). This is in line with previous findings that Fe oxidizers support higher diversity (24, 30). Differences in relative abundances of common lineages in microbial mats have been observed at sites with differing chemistry in previous studies (30, 85, 93), emphasizing the influence of vent fluids on microbial mat communities.

**Figure 7.**
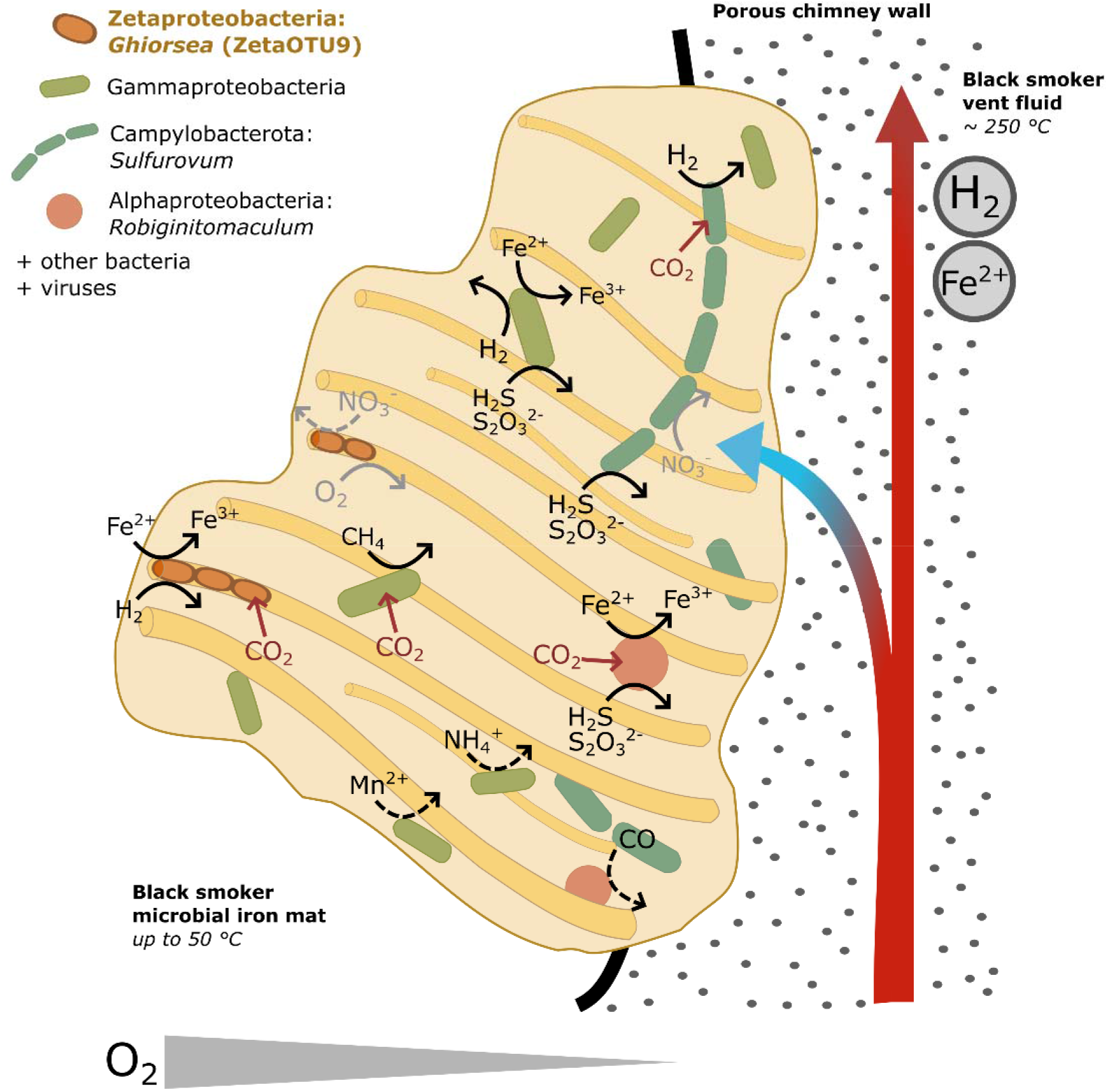
Conceptual model of Fe mats on black smoker chimneys at Fåvne vent field. Fe mats are found on black smoker chimneys with focused flow high-temperature venting of fluids containing iron and hydrogen. The temperature in the Fe mat close to the chimney exterior was ∼50 °C. The model is based on metabolic reconstruction of MAGs of the most abundant microbial groups.

The mixing of oxygen-rich seawater and reduced vent fluids in the porous chimney structures at Fåvne give rise to steep chemical gradients which is reflected in the available electron donors and acceptors utilized within the Fe mat (Figure 5). A strong association of Fe and Mn has been shown for hydrothermal fluids elsewhere (70) and at Fåvne genes likely involved in manganese oxidation or detoxification were detected in proteomics analysis (Figure 5, Table S10, Supplementary Material 1 Text2). Consistent with the notion that Fe(III) oxyhydroxides adsorb heavy metals (8), the presence of various heavy metal resistance genes in the Fe mat (Table S7) suggests adaptation to heavy metals.

### Diversity of Fe oxidation based on Cyc2 genes at Fåvne

Potential for iron oxidation at Fåvne vent field is not limited to Zetaproteobacteria based on the presence of Cyc2 genes across several phyla. The identification of *cyc2* in abundant members of the Gammaproteobacteria and Alphaproteobacteria may indicate a broader taxonomic range for iron oxidation (Figure 4, Figure 5), as seen in previous studies (34, 51, 52, 82, 94, 95). Based on metabolic profiling, MAGs possessing *cyc2* seem able to use oxygen and nitrate (Figure 5). The presence of nitrate and nitrite reductase genes in Zetaproteobacteria MAGs suggests a possibility of an advantageous metabolic plasticity in Zetaproteobacteria able to reduce nitrate and nitrite in the absence of oxygen. Such an anaerobic metabolism has not yet been observed in isolates under laboratory conditions (23, 77) and Zetaproteobacteria terminal oxidases are predicted to be highly expressed (Table S12). Similarly, in addition to the common microaerophilic Zetaproteobacteria, anaerobic iron oxidizers have been detected in deeper layers of Fe mats at Kama‘ehuakanaloa (Lō‘ihi) (4). It remains unknown whether the widespread occurrence of Cyc2 genes at Fåvne is involved in Fe oxidation to obtain energy for carbon fixation in several lineages or whether some of these microorganisms rather use the Fe oxidase in other processes such as detoxification (52, 96). It is however unlikely that Fe-oxidizing *Ghiorsea* at Fåvne are competing with other organisms for Fe resources, as there is an abundant supply of Fe(II) in the venting fluids (55).

### Porous black smoker chimneys support growth of Fe mats with iron and hydrogen at Fåvne

Previous studies of Fe mats constructed by Zetaproteobacteria have generally focused on low-temperature diffuse-venting areas rather than high-temperature hydrothermal vents (4, 25, 38). Although Fe mats on black smoker chimneys have been observed (32), the communities and interactions of their members have not yet been characterized. Whereas Fe mats studied previously are associated with fluids depleted in H_2_ (22, 27, 30, 34), vent fluids at Fåvne contain both H_2_ and Fe(II) as abundant energy sources for Fe mats growing on black smoker chimney surfaces (55). Other hydrothermal vents with similar geochemistry to Fåvne, such as Rainbow at Mid-Atlantic Ridge and Beebe’s vents (Piccard) at Mid-Cayman Rise where both H_2_ and Fe(II) are present, have chimneys that contain much higher temperature mineralized conduits that are likely not as porous (97, 98), preventing the establishment of a chemical gradient at the chimney surface and possibly unable to sustain large Fe mats. In contrast, the chimneys at Fåvne appear to be highly porous, visibly allowing vent fluid to permeate through to the chimney surface. This fluid flow creates a suitable environment for microbial life to access electron donors at warmer temperatures and form dense Fe mats.

## 4 Conclusion

The presence of Fe(II) and H_2_ in the fluids at the Fåvne vent field offers a unique opportunity to investigate the interactions and adaptations of Fe oxidizers in response to the presence of abundant H_2_, providing valuable insights into their physiology and ecological dynamics. Our study is a first look into the microbial communities of black smoker Fe mats and the first microbiological exploration of the newly discovered Fåvne hydrothermal vent field. The findings strongly suggest that Zetaproteobacteria of Fe- and H_2_-oxidizing genus *Ghiorsea* at Fåvne produce Fe(III) oxyhydroxide sheaths and form dense Fe mats. With these Fe(III) oxyhydroxide structures, *Ghiorsea* provide the environment for other microorganisms, ultimately maintaining the carbon, nitrogen, sulfur, and iron cycling in the Fe mats. Exclusive presence of Fe(II)- and H_2_-oxidizing *Ghiorsea* in the black smoker Fe mat exposed to abundant H_2_ compared with occupation by diverse Zetaproteobacteria without hydrogenases at likely low H_2_ environments at Fåvne supports the notion that H_2_ availability plays a crucial role in driving the niche partitioning of Zetaproteobacteria.

## 5 Material and Method

### 5.0 Sampling site

The Fåvne vent field is located at 72°45.4’ N, 3°49.9’ E on the Mohns Ridge section of the series of Arctic Mid-Ocean Ridges (AMOR) at 3030 m below sea level (54–56) (Figure S1). Black smoker chimneys have a porous structure and are rich in iron oxide and oxyhydroxide minerals, with high cobalt concentration of hydrothermal deposits (54, 56). The black smoker hydrothermal fluids there are characterized by abundant iron and hydrogen (55).

### 5.1 Sample collection

Iron microbial mat samples were collected using an Ægir6000 remotely operating vehicle (ROV) on board the R/V G.O. SARS in June 2019, equipped with a biosyringe (a hydraulic sampling cylinder) connected to the ROV manipulator arm (Figure 1a). Temperature (±1 °C uncertainty) in the iron microbial mat were taken in real-time using a temperature probe attached to the isobaric gas-tight fluid sampler snorkel inlet (99), which was used for vent fluid sample collection (55). Comprehensive vent fluid chemical compositions will be presented in a separate publication. Iron microbial mat (FeMat) was collected on the exterior of a 13 m tall active black smoker chimney, 1-2 m below the orifice of the vent, on the North Tower vent at coordinates 72.7567, 3.8336 and at 3025 m depth (Figure 1, Supplementary Material 3, Supplementary Material 4 Table S1). Additional samples of the iron microbial mat mixed with underlying chimney, chimney, and iron oxide deposits were collected (Supplementary Material 4 Table S1). Samples retrieved were centrifuged at 6000 rcf for 5 minutes and the supernatant was removed. Iron microbial mat pellet for on-ship metagenome sequencing using Nanopore MinION was processed directly. Aliquots for other analyses were frozen in liquid nitrogen and stored at −80 °C until processing. Samples for scanning electron microscopy were fixed in 2.5% glutaraldehyde and stored at 4 °C until further processing.

### 5.2 Scanning electron microscopy (SEM) and elemental composition analysis

Fixed samples for SEM were filtered onto 0.2 μm polycarbonate filters with subsequent incubation in a series of increasing ethanol concentrations to remove water with and without critical point drying in CO_2_. Samples were then sputter coated with Au/Pd or Ir. SEM images were produced using a Zeiss SUPRA 55VP scanning electron microscope at ELMILAB (Laboratory for analytical electron microscopy) at the Department of Earth Science (Faculty of Mathematics and Natural Sciences, University of Bergen). For analysis of elemental composition with a scanning electron microscope, energy-dispersive X-ray spectroscopy (EDS) and Pathfinder software were used at the same laboratory.

### 5.3 Fluorescence *in situ* hybridization (FISH)

Samples were fixed in 2% formaldehyde at 4 °C overnight, spread on microscopy slides, air-dried, and embedded in 0.5% low melting point agarose. For visualizing Zetaproteobacteria, the Zeta674 probe labelled with Atto488 fluorochrome was used (22). The Zeta674 probe specificity was analyzed and the probe was successfully hybridized *in silico* using the SILVA Test-Probe tool, local BLAST and the 16S rRNA gene sequence of the reconstructed *Ghiorsea* MAG (Faavne_M6_B18). FISH was performed according to a previously published protocol (100). Slides were incubated at 46 °C for 1 h with 20-30% formamide hybridization buffer in a hybridization chamber. The probe was added, followed by hybridization for 2 h at 46 °C. Slides were then incubated in a washing solution (0.1 M NaCl, 20 mM Tris-HCl (pH 8.0), 5 mM EDTA, and 0.01% SDS) at 48 °C for 15 minutes, washed twice with PBS and airdried. Vectashield antifade solution was added. Slides were visualized with fluorescence microscopy using an overlay of phase-contrast and florescence images. SYBRGreen was used to visualize all cells.

### 5.4 Genome-resolved metagenomics

#### 5.4.1 On-ship Nanopore MinION sequencing workflow

DNA extraction, sequencing and preliminary analysis was performed on board the research vessel during the expedition. DNA was extracted from a 1 ml FeMat sample aliquot using FastDNA Spin kit for soil (MP Biomedicals) according to the manufacturer’s protocol. Metagenomic sequencing of total DNA was carried out using the rapid sequencing library (SQK-RAD004) and the Oxford Nanopore Technologies MinION 1Mk1B sequencer equipped with a FLO-MIN106 SpotON Flow cell v. R9. Sequencing and raw data acquisition was controlled with the MinKNOW software. Basecalling was performed with a local version of the guppy basecaller v. 3.4.4 (https://community.nanoporetech.com). Filtering of raw reads on length and quality was performed twice using Nanofilt v.2.5.0 as part of the NanoPack (101) and Porechop v.0.2.4 (https://github.com/rrwick/Porechop) (sequencing and filtering statistics in Table S1A and S1B).

#### 5.4.2 llumina sequencing workflow

Whole-sample genomic DNA was extracted using MO BIO Power Soil kit and sent to the Norwegian Sequencing Centre (University of Oslo, Norway) for shotgun metagenomic sequencing. 150 bp paired-end sequencing was performed using an Illumina NovaSeq S4 flow cell. Raw reads were scanned for quality, duplication rate, and adapter contamination using FastQC v0.11.9 (https://github.com/s-andrews/FastQC), and concurrent visualization of the reports across samples was carried out in MultiQC (102). Strand-specific quality filtering methods recommended (103) were implemented through use of the “iu-filter-quality-minoche” script of the illumina-utils python package (104). Quality-filtered reads were subsequently cleaned of contaminating human DNA by mapping reads to the hg19 human genome with a mask applied to highly conserved genomic regions using the bbmap.sh script within the BBTools package (105) and human genome mask developed by Bushnell.

Sequence reads were assembled by individual metagenomic sample with MEGAHIT v1.2.9 (106) using a minimum contig length of 1000 bp. Reads from each sample were consecutively mapped to individual Illumina sample assemblies, effectively “co-mapping”, using Bowtie2 v.2.4.2 (107) then subsequent indexing with Samtools v.1.11 (108). Binning and quality procedures were identical to those carried out as detailed for MinION sequencing with the exception of inclusion of MaxBin2 v 2.2.4 as an additional binning software used before implementation of DASTool. File manipulation, contig database creation and profiles were accomplished with scripts from the Anvi’o v.7 platform (109).

##### Metagenome assembly strategy

Assembly of the ONT filtered reads was performed using the wtdbg2 v.2.5 long-read assembler (*options:* -p 21 -AS 2 -s 0.05 -L 2500--edge-min 2 --rescue-low-cov-edges) (110). Sequential polishing of the initial assembly was conducted twice with Racon v.1.4.3 (111) and Medaka v.0.8.2 (https://github.com/nanoporetech/medaka). Reconstruction of metagenome-assembled genomes (MAGs) was performed in a combination using CONCOCT (112), MetaBAT (113, 114) and DASTool (115).

Hybrid assembly of Nanopore and Illumina reads was also performed using metaSPAdes (116). This hybrid assembly had lower quality than wtdbg2 and MetaFlye-only and Illumina polished MetaFlye assemblies (Table S2), with lower quality bins and fewer 16S rRNA genes assigned to the genomes. Nanopore-only-based assembly generated 1 Zetaproteobacteria MAG in FeMat sample, while 2 distinct Zetaproteobacteria genomes were obtained using polished MetaFlye assembly. With this in mind, we decided to go forward with the MetaFlye long read assembly polished with Illumina reads. Assembly read information and quality metrics are shown in Table S2. Several other assembly and binning strategies were attempted and compared using QUAST v.5.0.2 with MetaQUAST output (117).

#### 5.4.3 Combining long reads and short reads

To obtain high quality metagenome-assembled genomes (MAGs) with better sequencing depth, a MetaFlye long-read assembly, done with Flye v2.9 and filtered nanopore reads (118), was polished using Illumina short reads using Pilon v1.23 (119). Illumina short reads were mapped to the assembly using bwa v.0.7.17 (120) and minimap2 for Nanopore long reads (121). A mapping file was then reformatted using samtools (108). Automatic MAG reconstruction was performed using metaWRAP v.1.3 (122) which implements the combinatorial use of MaxBin2 (123), CONCOCT (112) and MetaBat2 (114). MAGs were manually refined using the Anvi’o v.7 platform (109). Quality and completeness of individual MAGs were assessed on the presence of lineage-specific, conserved single-copy marker genes using CheckM v1.0.7 (124) and using CheckM2 v1.0.2. (125).

MAGs generated using MetaFlye, Pilon and metaWRAP and the ones generated using only Illumina reads, were dereplicated at 98% ANI using dRep v3.2.2 (126). These included all MAGs with at least 50% completeness and maximum 10% redundancy and all MAGs that had at least 0.5 coverage in the iron microbial mat metagenome (FeMat sample; 108 MEGAHIT Illumina MAGs and 19 MetaFlye, Pilon and metaWRAP MAGs). The dereplication resulted in 111 MAGs. Relative abundances were calculated using the abundance output of relative coverage within one sample (Anvi’o v.7), and this was normalized to one.

### 5.5 Taxonomic classification

The reconstructed MAGs were taxonomically classified using the genome taxonomy database tool kit gtdbtk v.1.7.0 (127) using the database GTDB 202 release. In addition, ZetaHunter v1.0.11 was used for assigning taxonomy to the 16S rRNA gene of the Zetaproteobacteria MAGs (42), classifying sequences into Zetaproteobacteria operational taxonomic units (ZetaOTUs) at 97% similarity. Based on ZetaHunter cut-offs, we excluded all ZetaOTU classifications below 75% entropy. An overall taxonomic classification of Illumina metagenomic reads was performed with PhyloFLASH v.3.4 (128) based on 16S rRNA genes using SILVA release 138 taxonomy as reference.

### 5.6 Genome database of Zetaproteobacteria

Zetaproteobacteria MAGs were reconstructed from samples of iron microbial mats, a chimney and iron deposit at Fåvne (Supplementary Material 4 Table S1). The choice was made to concentrate efforts on the black smoker iron microbial mat (FeMat) after identifying the presence of only the genus *Ghiorsea* and iron oxide sheaths and since FeMat was the most precise sample of the iron microbial mat. All publicly available Zetaproteobacteria genomes (73; taxid 580370) and corresponding metadata at NCBI GenBank were downloaded using ncbi-genome-download v.0.3.0 (https://github.com/kblin/ncbi-genome-download/) on the 19th of October 2021. In addition, publicly available genomes of Zetaproteobacteria were downloaded from Genomes from Earth’s Microbiome (129), Joint Genome Institute Integrated Microbial Genomes (JGI IMG), and from public repositories stated in selected studies (4, 34, 65, 130). Additional genomes of Campylobacterota, Gammaproteobacteria and Alphaproteobacteria closely related to the Fåvne MAGs were downloaded from NCBI as references. A threshold cut-off of high and medium quality genomes (min. 50% completeness, max. 10% redundancy) was used before further analysis. Phylogenomic analyses included 148 Zetaprotobacteria genomes in addition to the MAGs from this study. All selected genomes are presented in Supplementary Table S4.

The combination of average nucleotide identity (ANI), average amino acid identity (AAI) and alignment fraction (AF) provides an objective measure of genetic relatedness between Zetaproteobacterial genomes. The proposed species cut-off at ∼95% ANI (63, 64), ∼95 to 96% AAI (131) and 60% AF (132) was used, with the genus boundary at 65% AAI (133). ANI analysis based on the BLAST algorithm (ANIb) was performed using the anvi-compute-genome-similarity program within Anvio v.7.0 (109); --program pyANI --method ANIb (https://github.com/widdowquinn/pyani). AAI analysis was performed using ezAAI (134). AF was calculated using FastANI within Anvio. The graphical heatmap and dendrogram of percentage identities were plotted using gplots package in R.

Optimal growth temperatures were predicted for Zetaproteobacteria MAGs (Figure S9) using genomic features and regression models (135) (https://github.com/DavidBSauer/OGT_prediction). The models employed were Superkingdom Bacteria regression models that take into consideration the common absence of 16S rRNA gene and genome incompleteness in MAGs.

### 5.7 Phylogenetic and phylogenomic analyses of Zetaproteobacteria

Phylogenomics was used to determine the closest evolutionary relationships. Single copy marker genes present in all genomes were detected and extracted using Anvio v.7.0 (109) with anvi-get-sequences-for-hmm-hits, using Anvio’s Bacteria_71 and GTDB’s bac_120 collection of single copy marker genes (127). Selection of marker genes was based on genes being present only in a single copy, found in at least 70% of all Zetaproteobacteria genomes and supporting Zetaproteobacteria monophyly in individual marker phylogenetic trees (Table S5). Selected marker genes were comparable to the amount of single copy genes used for evolutionary placement of diversity within Zetaproteobacteria previously (1). Single-copy marker genes were manually checked and phylogenetic trees of selected individual protein sequences constructed using ultrafast bootstrapping (136). Selected individual marker gene alignments were constructed using MAFFT L-INS-i v7.397 (137), trimmed with trimAl v1.4. rev15 with selected parameters -gt 0.5 -cons 60 (138) and concatenated using catfasta2phyml (https://github.com/nylander/catfasta2phyml). A maximum-likelihood tree was constructed with IQ-TREE v2.0.3 (139) with non-parametric bootstrapping and using the best-fit model Q.pfam+F+I+I+R7 as determined by ModelFinder (140).

16S rRNA gene sequences were extracted using barrnap v.0.9 (https://github.com/tseemann/barrnap, settings: --kingdom ‘bac’ --evalue 1e-20) and only sequences longer than 500 bp were kept. The alignment was constructed using MAFFT L-INS-i v7.397 (137), manually inspected for non-matching sections and trimmed with trimAl v1.4. rev15 with trimming option -gappyout (138). Using sequences of comparable length, a maximum-likelihood tree of 16S rRNA genes was constructed using IQ-TREE v2.0.3 (139) with non-parametric bootstrapping and the best-fit model GTR+F+I+I+R3 as determined by ModelFinder (140).

### 5.8 Functional annotation and genome comparison

Gene calling and functional annotation of MAGs was performed with an automated pipeline (141) conducting separate searches against Prokka v1.14 (142), NCBI COG (downloaded from NCBI webserver in February 2021), arCOG (version from 2018) (143), KEGG (downloaded in February 2021) (144), Pfam (release 33.0) (145), TIGRFAM (release 15.0) (146), CAZy (dbCAN v9) (147), Transporter Classification Database (downloaded from TCDB webserver in February 2021) (148), HydDB (downloaded from HydDB webserver in February 2021) (67) and NCBI_nr (downloaded from NCBI webserver in February 2021). Genes of interest (presence/absence) were determined for metabolic reconstruction mainly based on KEGG and TIGRFAM annotations (Table S7A), with the main functions discussed in the article manually inspected. Genes for CO_2_ fixation pathways were screened using a customized script based on KEGG decoder v1.2.1 (149, 150). Iron oxidation genes were identified using FeGenie v1.1 tool (151) and manganese oxidation genes were annotated using MagicLamp v1.0 with curated lithotrophy HMMs (152). Potential stalk formation gene homologues were identified based on a local BLAST together with previously studied sequences, % identity cut-offs and average gene lengths previously defined (46). Metal resistance genes were identified in metagenome-resolved genomes and assembly using the BacMet database (version 2.0) with experimentally confirmed and predicted resistance genes. Predicted resistance gene identification criterion of min. 85% sequence identity with the full-length coverage of the short reads (75-300+ bps) was used as advised by the database authors.

### 5.9 Codon bias expression prediction

Codon bias gene expression levels was predicted using coRdon R package (v 1.8.0, https://github.com/BioinfoHR/coRdon), based on the MILC (Measure Independent of Length and Composition) and MELP (MILC-based Expression Level Predictor) values (153).

### 5.10 Phylogenetic tree of Cyc2

Cyc2 sequences were identified in MAGs present in the black smoker iron microbial mat and in all Zetaproteobacteria from all sampled Fåvne sites using FeGenie (151). Additional Cyc2 identifications were made from the top 10 hits using a blastp alignment based on the GenBank and NCBI_nr database. References were downloaded with BatchEntrez and reannotated as Cyc2 using FeGenie. Sequences shorter than 300 and longer than 600 amino acids were filtered out. Identical sequences were dereplicated using clustering with CD-HIT v4.8.1 (154). The resulting sequences of similar lengths were aligned using MAFFT L-INS-I v7.397 (137), manually checked in AliView v1.26 (155), and trimmed using trimAl v1.4. rev15 with -gt 0.7 (positions in the alignment with gaps in 30% or more of the sequences were removed) (138). A maximum likelihood phylogenetic tree was constructed with IQ-TREE v2.0.3 (139) using an alignment of 115 sequences with 388 positions and the best-fit model Q.pfam+F+I+I+R5 according to ModelFinder (140). Branch support values were calculated with standard bootstrapping with 1000 iterations. The tree was rooted at midpoint.

### 5.11 Phylogenetic tree of Ni,Fe hydrogenase

The protein sequences of Ni,Fe large subunit hydrogenase 1d were identified from several sources: from the Fåvne iron microbial mat MAGs, the reference Zetaproteobacteria, along with their closest relatives. Closest relatives were identified by BLAST Diamond annotations using HydDB (downloaded from HydDB webserver in February 2021) (67), the top 50 hits using an additional blastp alignment using GenBank and nr database, and blastp using a JGI IMG database gene search at 85% identity threshold (August 2022). 213 NiFe large subunit hydrogenase 1d reference sequences from HydDB were added (downloaded from HydDB webserver in March 2021). Sequences shorter than 460 amino acids were filtered out. The resulting sequences of similar lengths were aligned using MAFFT L-INS-i v7.397 (137), manually checked in AliView v1.26 (155) and trimmed using trimAl v1.4. rev15 with -gt 0.5 -cons 60. A phylogenetic tree was constructed with IQ-TREE v2.0.3 (139) using an alignment of 317 sequences with 595 positions, based on maximum likelihood and the best-fit model LG+I+I+R7 according to ModelFinder (140). Branch support values were calculated with standard bootstrapping with 1000 iterations. Redundant sequences from several sources (NCBI, IMG JGI) were pruned, leaving only one sequence representative. The tree was rooted at midpoint. Environment data was pulled from available metadata and taxonomy from NCBI with corrections based on GTDB where genomes were present.

### 5.12 Viral genomes

CheckV v.0.8.1 (156), VIBRANT v.1.2.1 (157) and DeepVirFinder (158) were used to search for viruses in the unbinned sequences. Quality of the viral genomes was checked using CheckV and reported according to minimum information requirements on uncultivated viral genomes (159). We predicted host-virus associations in the iron microbial mat using host CRISPR-spacers, integrated prophage, host tRNA genes, and k-mer signatures with VirMatcher, accessed in September 2022 (https://bitbucket.org/MAVERICLab/virmatcher/). This was done using minCED (v. 0.4.2; https://github.com/ctSkennerton/minced), BLAST (160), promiscuous tRNA sequences (161), tRNAscan (162) and WIsH (163). Score of 3 was used as a threshold value to assign hosts based on previous approaches (164). Host-virus pairs were analyzed also with PHP host predictor software using K-mer predictions (165).

## 6 Data availability

All MAGs in the study were deposited in NCBI and accession numbers with associated BioProject and BioSamples with corresponding metadata are listed in Supplementary Material 4 Table S1 and Supplementary Material 2 Table S4 and Table S5. Supplementary material is available online, deposited in a Zenodo repository at https://doi.org/10.5281/zenodo.8058699. Code used for the analyses is available at https://github.com/MicrobesGonnaMicrobe/Faavne_IronMats_Analysis.

## 7 Supplementary Material

Supplemental material is available online, deposited in a Zenodo repository at https://doi.org/10.5281/zenodo.8058699.

**Supplementary Material 1.** Supplementary Data and Figures.

**Supplementary Material 2.** Supplementary Tables.

**Supplementary Material 3.** Sampling video of black smoker iron microbial mats at Fåvne hydrothermal vent (North Tower vent site).

**Supplementary Material 4.** Sampling locations.

## 9 Author Contributions

Author contributions are assigned using CRediT roles.

**Petra Hribovšek**: Conceptualization (lead), data curation (equal), formal analysis (lead), investigation (lead), methodology (lead), visualization (lead), writing – original draft (lead), writing – review & editing (lead). **Emily Olesin Denny**: Data curation (lead), writing – original draft (supporting). **Håkon Dahle**: Writing – original draft (supporting). **Achim Mall**: Data curation (equal), software (lead), writing – original draft (supporting). **Thomas Øfstegaard Viflot**: Formal analysis (supporting), investigation (supporting). **Chanakan Boonnawa**: Formal analysis (supporting), investigation (supporting). **Eoghan P. Reeves**: Formal analysis (supporting), investigation (supporting), writing – original draft (supporting), writing – review & editing (supporting). **Ida Helene Steen**: Conceptualization (lead), formal analysis (supporting), funding acquisition (lead), investigation (supporting), methodology (supporting), supervision (lead), writing – original draft (supporting), writing – review & editing (supporting). **Runar Stokke**: Conceptualization (lead), data curation (lead), formal analysis (supporting), funding acquisition (lead), investigation (supporting), methodology (lead), supervision (lead), project administration (lead), writing – original draft (supporting), writing – review & editing (supporting).

## 11 Acknowledgments

We thank the ROV Ægir6000 operator team and the R/V G.O. SARS crew for assistance during sampling, and Steffen L. Jørgensen for organizing the 2019 research cruise to the Fåvne vent field. We thank Irene Heggstad for assistance with scanning electron microscopy. The computations associated to taxonomic classification and annotation of genes were performed on resources provided by Sigma2 - the National Infrastructure for High Performance Computing and Data Storage in Norway.

